# Peripheral membrane proteins modulate stress tolerance by safeguarding cellulose synthases

**DOI:** 10.1101/2022.05.24.493199

**Authors:** Christopher Kesten, Álvaro García-Moreno, Vítor Amorim-Silva, Alexandra Menna, Araceli G. Castillo, Francisco Percio, Laia Armengot, Noemi Ruiz-Lopez, Yvon Jaillais, Clara Sánchez-Rodríguez, Miguel A Botella

## Abstract

Controlled primary cell wall remodeling adapts plant growth under stressful conditions, but how these changes are conveyed to adjust cellulose synthesis is unknown. Here, we identify the Tetratricopeptide Thioredoxin-Like (TTL) proteins as new members of the cellulose synthase complex (CSC) and describe their unique and hitherto unknown dynamic association with the CSC under cellulose-deficient conditions. We find that TTLs are essential for maintaining cellulose synthesis under salinity stress, establishing a stress-resilient cortical microtubule array, and stabilizing CSCs at the plasma membrane. To fulfill these functions, TTLs interact with Cellulose Synthase1 (CESA1) and engage with cortical microtubules to promote their polymerization. We propose that TTLs function as bridges connecting stress perception with dynamic regulation of cellulose biosynthesis at the plasma membrane.

**One Sentence Summary:** TTLs are peripheral membrane proteins that maintain the integrity of the cellulose synthase complex upon adverse conditions.

## Main Text

Stress-sensing mechanisms enabling rapid, adaptive, and controlled remodeling of the primary cell wall mitigate perturbations on plant cellulose synthesis under environmental stress (*1, 2*), thereby alleviating biomass and growth reduction. Yet, how fine-tuning of the highly dynamic cellulose synthase (CESA) complex (CSC) is achieved under stress remains largely unknown. Various signaling pathways stimulate the recruitment of cytosolic Tetratricopeptide Thioredoxin-Like (TTL) proteins to the plasma membrane (*3*) where, acting as scaffolds, they amplify and stabilize the initial signals. *TTLs* were identified through forward genetic screens for mutants with reduced root growth and isotropic cell expansion under elevated NaCl concentrations. Since these are also common phenotypes of cellulose-deficient mutants (*4, 5*), we explored potential cellulose synthesis defects in *ttl* mutants under salt stress. *ttl1, ttl1ttl3*, and *ttl1ttl3ttl4* roots were significantly shorter than wild type (WT; Col-0) roots under high (200 mM) NaCl concentrations *ttl1ttl3*- and *ttl1ttl3ttl4*- also showed exacerbated isotropic cell expansion compared to WT- or *ttl1*-roots (fig.S1A-B). Since the phenotypes of the *ttl1ttl3* double mutant were only mildly enhanced in comparison to WT by adding the *ttl4* knockout, we focused on the *ttl1ttl3* for further characterization of the TTLs. Less crystalline cellulose accumulated in *ttl1ttl3* compared to WT roots and isotropic growth defects occurred in dark-grown hypocotyls from the former, but not the latter (fig. S1C-F), a phenotype replicated with the primary wall cellulose synthesis inhibitor isoxaben (*6*); fig.S1G-I). Collectively, these results link *TTL1* and *TTL3* to cellulose synthesis at the primary cell wall.

Since TTL functions are underpinned by their unique subcellular dynamics, we explored potential co-/re-localization between TTL1/3 and the CSC. The salt sensitivity of *ttl1ttl3ttl4* was rescued in the *ttl1ttl3ttl4 pTTL3:TTL3-GFP* line (fig. S1J-K) as was that of *ttl1ttl3* when complemented with *pTTL1:TTL1-GFP* (fig.S1L-M). Spinning-disk confocal microscopy in etiolated hypocotyls revealed a prominent cytoplasmic localization for TTL3-GFP and a weaker signal of motile foci at the plasma membrane, as confirmed by comparative immunoblot analyses between cytosolic and microsomal fractions (fig.S1N-O). TTL1-GFP from line *ttl1ttl3 pTTL1::TTL1-GFP* displayed a similar dual-localization but mainly in the roots elongation zone (fig.S1P) contrasting with TTL3-GFP broad expression in roots and hypocotyls (*3*). The motile plasma membrane-localized TTL1/3-GFP foci evoked those formed by CESAs (*7*), prompting us to test if TTLs indeed track together with them. In a dually-labeled *ttl1ttl3* line expressing *TTL3-GFP* and tdTomato (*tdT*)-*CESA6* (*8*), the motile TTL3-GFP foci were substantially obscured by strong cytosolic signals (Fig.1A). Use of an in-house image modification workflow (see *Methods*) to extract the foci from the cytosolic signal (Fig.1B, Movie S1) revealed that TTL3-GFP colocalizes and co-migrates with tdT-CESA6 at the plasma membrane (Fig.1B-D, Movie S1). Agreeing with their co-localization, TTL3-GFP and tdT-CESA6 moved bi-directionally with similar speeds (271 ± 86 nm/min, and 267 ± 93, respectively; Fig.1E).

**Fig. 1.**
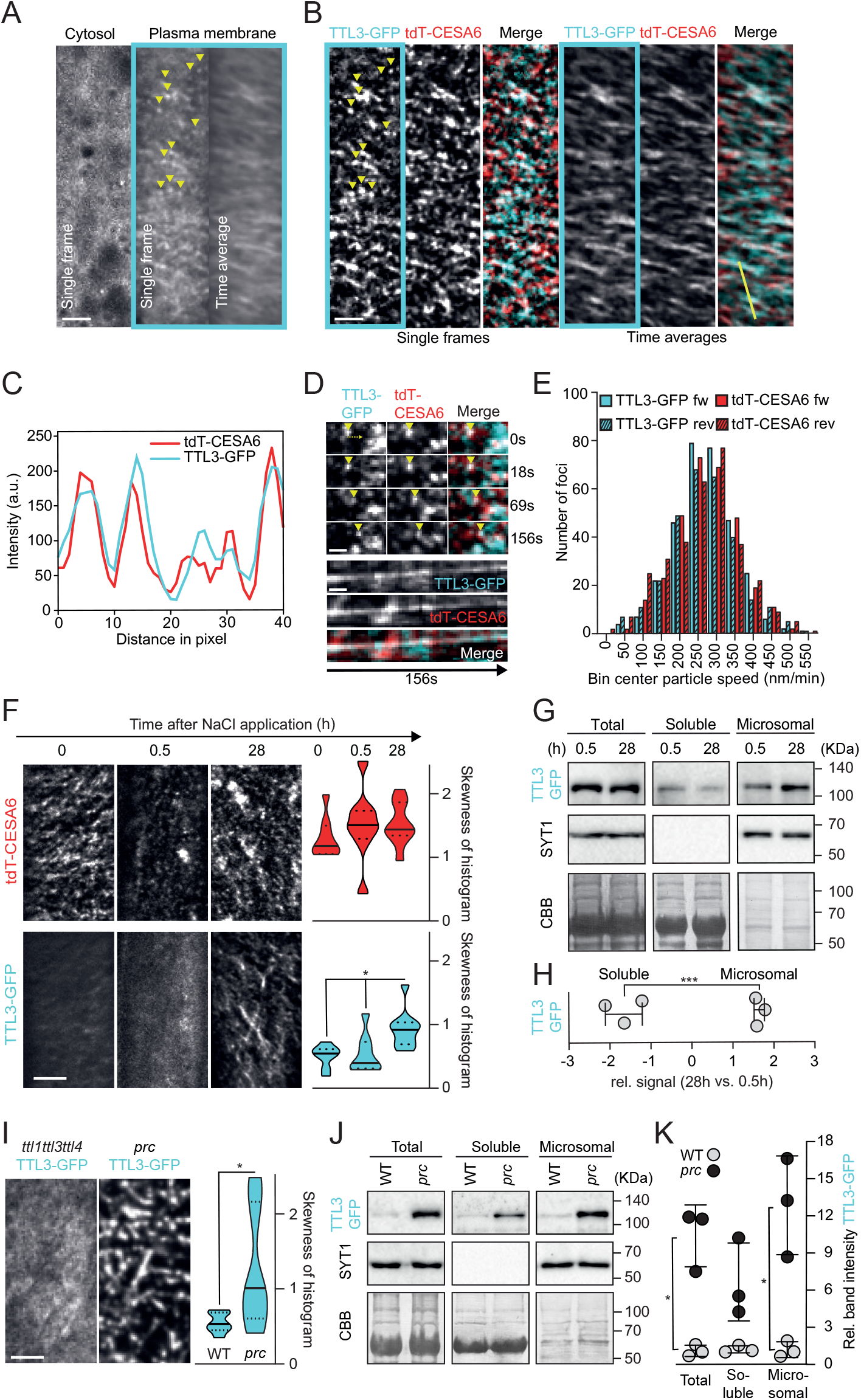
Salt stress and low cellulose content regulate TTL3 localization and dynamics. (**A)** TTL3-GFP in etiolated hypocotyl epidermal cells of *ttl1ttl3 TTL3-GFP tdT-CESA6* seedlings. Single frames and time-average projections show that TTL3-GFP is cytosolic but also forms motile particles at the plasma membrane (framed in cyan, yellow arrows). Bar=2.5μm. (**B)** Plasma membrane focal plane of the cell as in (A). TTL3-GFP particles co-localize with tdT-CesA6 at the plasma membrane (single frames). Time-average projections of TTL3-GFP and tdT-CESA6 reveal fluorescent foci’s co-migration. Bar=2.5μm. **(C)** Intensity plot of TTL3-GFP and tdT-C6 along the yellow line shown in (B). (**D)** Time frames of individual TTL3-GFP and tdT-CESA6 foci from corresponding movies. Upon their formation (yellow arrowhead at time 0s), foci co-migrate for at least 156s. Bar=1μm. (**E)** Bi-directional motility and speed distribution of TTL3-GFP and tdT-CESA6 foci. Forward denotes movement from left-to-right; reverse from right-to-left in relation to the major axes of analyzed cells. (**F**) Left: TTL3-GFP and tdT-CESA6 coverage at the plasma membrane in 3-days-old *ttl1ttl3* hypocotyl epidermal cells prior (time 0) and after exposure to 200 mM NaCl for the indicated durations. Bar =5μm. Right: Quantification of histogram skewness of images as shown in left panels. Centerlines in violin plots: medians; dotted lines: 25^th^ and 75^th^ percentiles. Welch’s ANOVA between CESA time-points: *p*-value 0.50; Welch’s ANOVA between TTL3-GFP timepoints: *p*-value ≤0.01. Dunnett’s T3 multiple comparisons: \**p*-value≤0.05. n≥18 cells from >8 seedlings per time-point in three independent experiments. (**G**) Total, soluble, and microsomal fractions were isolated from 3-days-old *ttl1ttl3ttl4* TTL3-GFP etiolated seedlings treated with 200 mM NaCl for 0.5h or 28h. TTL3-GFP detected with an anti-GFP antibody, control microsomal detected with an anti-SYT1 antibody. Coomassie brilliant blue (CBB) staining provides a loading control. (**H**) TTL3-GFP signal quantification from immunoblots as in (G). Band intensity is expressed as fold change of 28h vs 0.5h of 200 mM NaCl treatment in soluble and microsomal fractions. N=3 independent experiments. Values are individual replicates ±SEM; Unpaired t-test; *p-value≤0.05. (**I**) Left: TTL3-GFP coverage at the plasma membrane in 3-day-old *ttl1ttl3* or *ttl1ttl3prc1-1* etiolated seedlings. Bar=5μm. Right: Histogram skewness of images as shown in left panels. Violin plots: as in (F); n =13 cells from at least eight seedlings from three independent experiments. Welch’s unpaired t-test; \**p*-value ≤0.05. (**J**) Fractionation analysis of TTL3-GFP in WT and *prc1-1* as in (G) but without salt treatment. (**K**) TTL3-GFP signal quantification from immunoblots as in (J). Band intensity data is expressed as relative band intensity in total, soluble and microsomal subcellular fractions. The TTL3-GFP immunoblot signal intensity was normalized to that of SYT1 for the input and microsomal fractions, and to the CBB for soluble fractions. N=3 independent experiments. Values are individual replicates ±SEM; Welch’s unpaired t-test; \**p*-value≤0.05.

Given the *ttl* mutants hypersensitivity to NaCl, we investigated TTL3 localization and dynamics under control conditions or at 0,5h and 28h post-NaCl treatment, time-points at which CESA removal and subsequent plasma membrane re-population are optimally observed (*1*). In etiolated hypocotyls of *ttl1ttl3 TTL3-GFP tdT-CESA6* plants, TTL3-GFP was predominantly cytosolic in control conditions and 0,5h post-NaCl treatment; it was, however, strongly relocalized to tdT-CESA6-rich foci at the plasma membrane 28h post-treatment, with a concomitant depletion of the cytosolic signal (Fig.1F, see *Methods* for quantification). Furthermore, TTL3-GFP was significantly depleted from the soluble-, and enriched in the microsomal fraction 28h posttreatment, unlike the control microsomal protein SYT1 (*9*) (Fig.1G-H). Thus, TTL3 is a peripheral membrane protein that selectively re-localizes to the CSC upon stress. When introduced into *ttl1ttl3*, a null mutation in *CESA6* (*prc1-1*; (*10*)) aggravated the *ttl1ttl3* developmental defects, supporting genetic interactions between cellulose synthesis/content and TTL1/3 functions (fig.S2A-G). Accordingly, TTL3-GFP over-accumulated and formed predominantly motile clusters at the plasma membrane in *prc1-1* compared to *ttl1ttl3ttl4* plants, even in non-stressed conditions (Fig.1I). Furthermore, TTL3-GFP levels were increased by respectively ~6-versus ~12-folds in the cytosolic versus microsomal fractions of *prc1-1*- compared to WT-plants (fig.2 A-G).

**Fig. 2.**
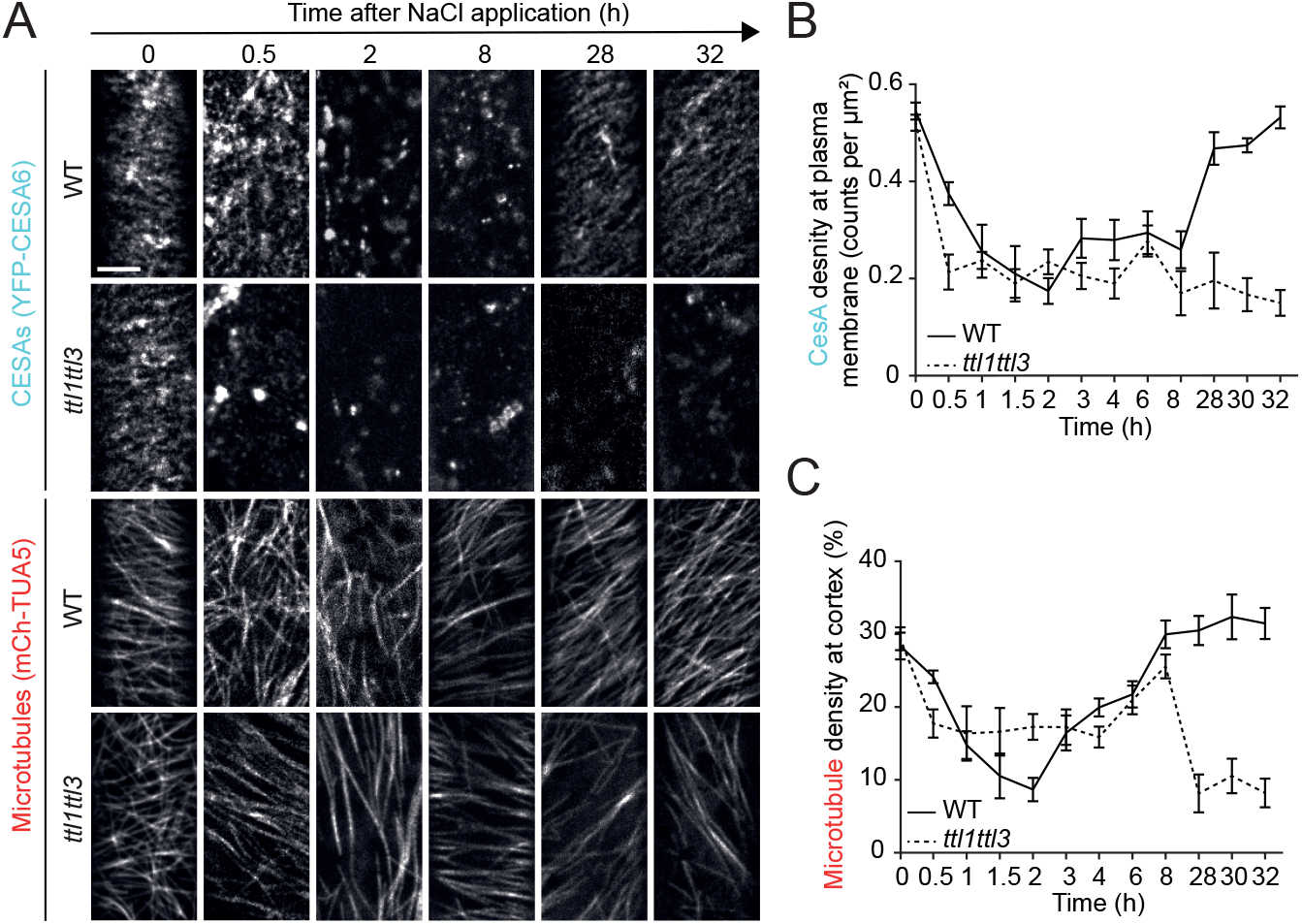
TTL3 regulates CESA activity and cortical microtubule polymerization under salt stress. (**A**) Microtubule and CESA coverage in WT and *ttl1ttl3* plants co-expressing mCh-TUA5 and YFP-CESA6 following exposure to 200 mM NaCl at the indicated time points. Time 0 shows cells just prior to NaCl treatment. Bar=5μm. (**B**) CESA density at the plasma membrane quantified from images as in (A). Time indicates the time after salt exposure. Two-way ANOVA analysis of CESA density; *p*≤ 0.001 (genotype), *p*≤ 0.001 (time), *p*≤ 0.001 (genotype × time). n≥27 cells from at least three seedlings per time point and three independent experiments. Values are mean ±SEM. (**C**) Microtubule coverage at the cell cortex quantified from images as in (A). Two-way ANOVA analysis of microtubule coverage is exactly as described for CESA density in (B).

To explore if TTL1/3 loss-of-activity might alter CESA removal and subsequent plasma membrane re-population under salt stress, we generated a *ttl1ttl3 YFP-CESA6 mCh-TUA5* (encoding *TUBULIN ALPHA-5*; (*11*)) line where imaged CESAs and microtubules in etiolated hypocotyls between 30min and 32h post-NaCl treatment. As reported, the microtubule array in WT plants depolymerized 2h post-treatment and reassembled within 8h thereof, in line with the decreased CESA6 density at the plasma membrane within 2h and its subsequent recovery within 28h (Fig.2A-C) (*1, 2, 12*). In contrast, cortical microtubules showed incomplete depolymerization 2h post-treatment in *ttl1ttl3* plants and, following a plateau-phase of ~8h, failed to reassemble (Fig.2A-C). Concurrently, a rapid decrease in CESA density at the plasma membrane within 30min post-treatment was never followed by re-population over the experiment’s time-course (Fig.2A-B). YFP-CESA fluorescence’s recovery after photobleaching was similar over time in WT and *ttl1ttl3* plants, ruling out a TTL3-dependent CESA6 delivery to the plasma membrane (fig.S2H) and suggesting that TTLs stabilize cortical microtubules and the CSC at the plasma membrane during cell adjustment to salt stress.

We tested if TTL3 directly interacts with primary CESAs using the yeast two-hybrid system (Y2H). Being full-length TTL3 unstable in yeast (*3*), we employed three partial constructs as preys (Fig.3A): two N-Terminal fragments encompassing the intrinsically disordered region (TTL3IDR), the IDR and the first two TPRs (TTL3Δ4TPR), or a C-terminal region containing all six TPRs and TRLX motif (TTL3ΔIDR). As baits, we used the cytosolic N-terminal domains (CESA-N) or the cytosolic catalytic domains of CESA1, CESA3, or CESA6 (CESA-C1,3,6, respectively). TTL3Δ4TPR interacted exclusively with CESA1-C, and neither TTL3IDR nor TTL3-ΔIDR interacted with any CESA domain (Fig.3B), despite all constructs being expressed (fig.S3A). In *N.benthamiana* transient expression, full-length TTL3 co-immunoprecipitated with CESA1-C, similarly to what we found using Y2H where TTL1Δ4TPR interacted with CESA1-C but not CESA1-N (Fig.3C and S3B). Therefore, TTL1/3 bind to CESA1-C *via* the N-terminal IDR domain and two first TPRs.

**Fig. 3.**
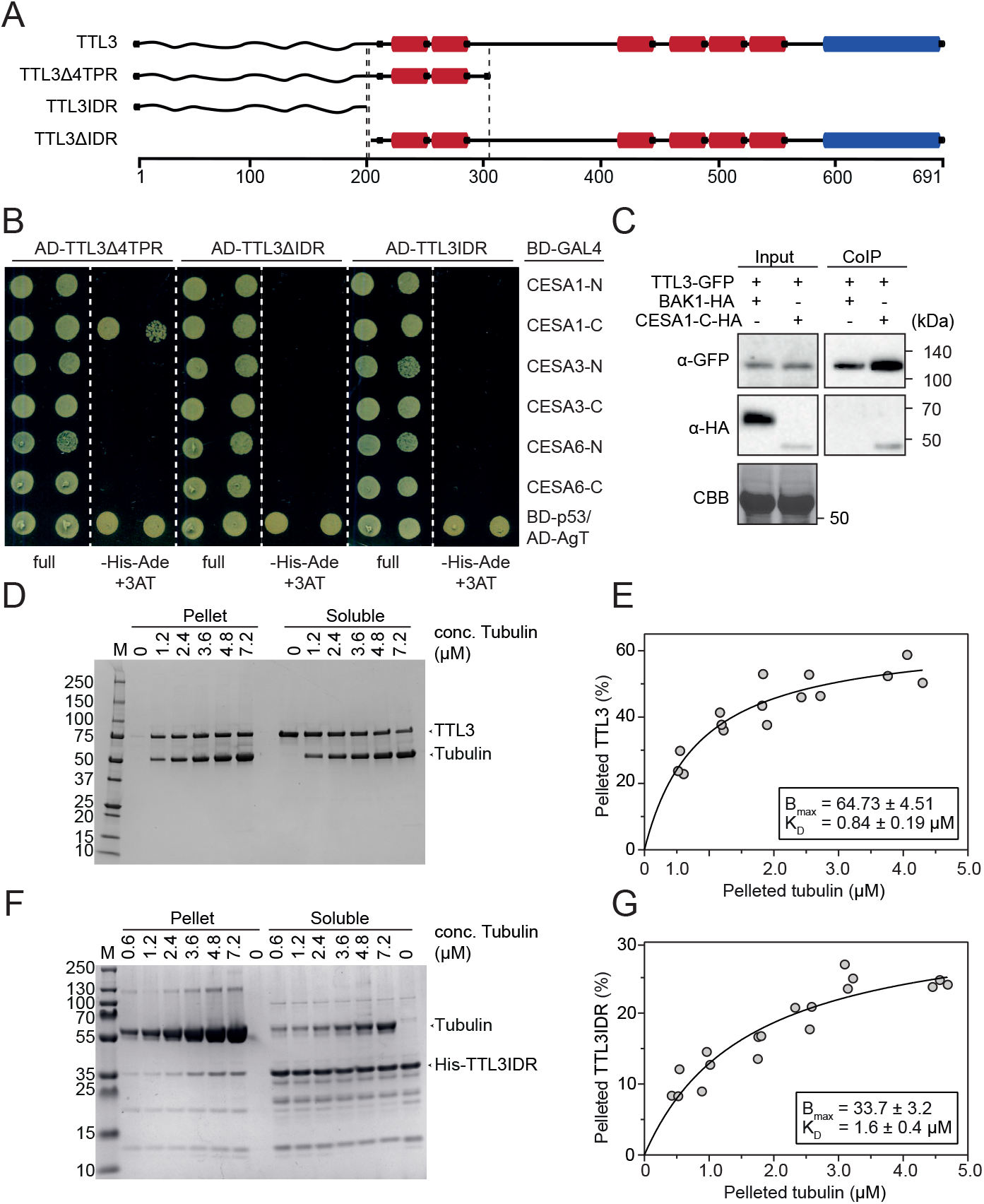
TTL3 is a microtubule-associated protein and interacts with the CESA1 catalytic domain. (**A)** Schematics of TTL3 and truncations thereof used here, with the predicted intrinsically disordered N-terminus (wavy line), 6 tetratricopeptide repeats (red), and one C-terminal thioredoxin-like domain (blue). (**B**) Yeast-two-hybrid assays of TTL3 fragments with the cytosolic CESA-N and CESA-C domains. BD-p53/AD-AgT was used as a positive control. Growth on non-selective media (full) and interaction-selective media (lacking histidine and adenine [–His–Ade] and supplemented with 3-amino-1,2,4-triazole [+3AT]) are shown. Photographs were taken after 3 days. Three independent experiments yielded similar results. (**C)** TTL3-GFP was transiently co-expressed with CESA1-C-HA (A) or BAK1-HA as a negative control in *N. benthamiana*. Total (input) and co-immunoprecipitated (CoIP) proteins were analyzed by immunoblotting using an anti-HA antibody. Equal loading was confirmed by Coomassie blue staining (CBB) of input samples. Two independent experiments yielded similar results. (**D**) Representative coomassie gel used to estimate the dissociation constants for TTL3 and microtubules. Tubulin and TTL3 molecular sizes are indicated with arrowheads. (**E**) The binding affinity of TTL3 to microtubules was determined by calculating their dissociation constant (K_D_) (0.84±0.19 μM; best-fit values ±Std. Error). TTL3 levels were kept constant while the amount of microtubules was increased. K_D_ was calculated by fitting a saturation binding curve onto the data, obtained from images as in (D); N=3 independent experiments. (**F**) Representative coomassie gel used to estimate the dissociation constant for 6xHis-TTL3IDR and microtubules. Tubulin and 6xHis-TTL3IDR molecular sizes are indicated by arrowheads. (**G**) The binding affinity of 6xHis-TTL3IDR to microtubules was determined by calculating their dissociation constant (K_D_) (1.6±0.4 μM; best-fit values ± Std. Error). 6xHis-TTL3IDR levels were kept constant while the amount of microtubules was increased. K_D_ was calculated by fitting a saturation binding curve onto the data, obtained from images as in (F); N=3 independent experiments.

The disturbed cortical microtubule recovery under NaCl stress in *ttl1ttl3* and high isoelectric point (>10) of its N-terminal IDR hinted at possible direct interactions between TTL3 and the negatively charged tubulin surface, which was tested with microtubule spin-down assays (*13*) using E. coli-expressed, full-length TTL3 (fig.S3C). Saturation binding experiments (*14*) revealed a dissociation constant (K_D_) of 0.84 ± 0.19 μM (Fig.3D-E), a microtubule-binding affinity within the range reported for CELLULOSE SYNTHASE INTERACTIVE 1 and Companion of Cellulose Synthase 1, known to interact directly with microtubules and CESAs (*1, 2, 15, 16*). A 6x-His-tagged version of the TTL3 IDR (TTL3IDR; fig S3D) also directly interacted with microtubules, albeit at lower affinity (K_D_ = 1.6 ± 0.4 μM; Fig.3F-G). To assess the role of TTL3 in microtubule polymerization, we performed microtubule turbidity assays. The TTL3IDR incurred an atypical polymerization curve with a short exponential phase followed by a linear increase (fig.S3E), contrasting taxol- and Microtubule Associated Protein-rich Fraction (MAPF)-based treatments leading to an exponential growth phase followed by a stationary phase of equal microtubule polymerization and depolymerization, as reported (*1*). To exclude that protein denaturation caused the linear phase in TTL3IDR-treated samples, pelleted reactions were analyzed by SDS-PAGE. Intact tubulin bands at 55 kDa could be observed in the TTL3IDR-, MAPF- and taxol-treated samples, while degradation products accumulated in both BSA- and buffer-treated samples (fig.S3F), indicating *bona fide* microtubule lattice stabilization by the TTL3IDR.

In summary, our results identify a novel family of proteins required for maintaining plant biomass production under stressful conditions. We show that TTL proteins are new CSC members that get dynamically recruited under stresses such as high salinity or reduced cellulose content to safeguard the cellulose synthesis machinery. Furthermore, TTL orthologs can be identified in all embryophytes down to the liverwort *Marchantia polymorpha*, the most basal lineage of extant plants (fig. S3G), while all other intrinsic components of the CSC emerged earlier in evolution (*17*). Thus, *TTL* genes are likely a key evolutionary acquisition that might allow plants to withstand stresses that are inherent to land colonization enabling a fine-tuned stabilization of the cellulose synthesis machinery.

## Acknowledgments

We are very grateful to Kamil Scklodowski for generating the CESAs constructs used for the Y2H assays and to Olivier Voinnet for critical reading of the manuscript. Live cell imaging was performed with equipment maintained by the Scientific Center for Optical and Electron Microscopy (ScopeM, ETH Zurich) and by the Center for Advanced Bioimaging (CAB) Denmark.

## Author contributions

MAB, CSR, CK, AGM, and VAS designed the research. CK conducted imaging and image analysis, heterologous protein expression, and microtubule-based assays. AGM and VAS generated the plant lines, physiological and the biochemical data. AM conducted cellulose measurements. AGC performed Y2H studies. FPV performed the physiological and the phylogenetic analysis. LA and YJ contributed to line generation. NRL contributed to biochemical analyses. Data analyses were led by CK, AGM, VAS, CSR, and MAB. CK, VAS, CSR, and MAB wrote the manuscript with major contribution from YJ. All authors read and provided feedback on the manuscript.

## Data and materials availability

All data is available in the manuscript or the supplementary materials.

## Supplementary Materials

### Materials and Methods

#### Plant material and growth conditions

*Arabidopsis thaliana* Columbia (Col-0) ecotype of was used in this study. The T-DNA insertions for TTL genes used in this study, have been described previously: *ttl1-2* mutant allele, in this study referred to as *ttl1* (AT1G53300) Salk_063943 (*1*); *ttl3* (AT2G42580) Sail_193_B05; *ttl4* (AT3G58620) Salk_026396; *ttl1ttl3* double and *ttl1ttl3ttl4* triple mutant (*2*). *prc1-1* single mutant was obtained from NASC and was described previously (*3*). The *ttl1ttl3ttl4* TTL3-GFP (*ttl1ttl3ttl4 pTTL3:TTL3g-GFP 2.4*) stable transgenic Arabidopsis line used in this study was described previously (*4*).The *ttl1ttl3* TTL1-GFP (*ttl1ttl3 pTTL1:TTL1g-GFP*) stable transgenic line were generated as described in “Plasmid constructs’’ and “Generation of transgenic plants’’. *ttl1ttl3* YFP-CESA6 mCherry-TUA5 was obtained by crossing *ttl1ttl3* with a YFP-CESA6 mCherry-TUA5 dual labeled line (*5, 6*), *ttl1ttl3* TdT-CESA6 TTL3-GFP was obtained by crossing *ttl1ttl3ttl4* TTL3-GFP with TdT-CESA6 (previously described in Sampathkumar et al., 2013), *prc1-1* TTL3-GFP was obtained by crossing *ttl1ttl3ttl4* TTL3-GFP with *prc1-1* and *prc1-1ttl1ttl3* was obtained by crossing *ttl1ttl3* with *prc1-1*. Seeds were surface sterilized using the chlorine gas method (*7*) and cold treated for 4 days at 4°C for stratification. For root elongation assays, seeds were sowed onto half-strength Murashige and Skoog (MS) agar solidified medium (1% [w/v] for vertical growth) containing 1.5% sucrose and subsequently grown under cool-white light (approx. 120 μmol photon m^-2^ s^-1^), at 22°C, with a long-day photoperiod (at 16 h light/8 h dark cycle). For etiolated seedlings analyses, seed were sowed onto half-strength MS agar solidified medium (1% [w/v] for vertical growth) containing 1% sucrose, stratified for 4 days, exposed to light (approx. 120 μmol photon m^-2^ s^-1^) for 3 h and subsequently grown in dark conditions at 22°C for 4 days. When required, seedlings were transferred to soil after 7 days of in vitro growth and watered every 2 days. In soil, plants were grown in a mixture of organic substrate and vermiculite (4:1[v/v]) at 22°C, with a long day photoperiod (approx. 120 μmol photon m^-2^ s^-1^).

#### Isoxaben and NaCl treatment

For *in vitro* assays, seeds were sowed in half-strength MS agar-solidified medium supplemented with 1% sucrose plus 1 nM Isoxaben (Santa Cruz Biotechnology) and 5 day-old etiolated seedlings were photographed and analysed for hypocotyl length and width. For salt stress, seedlings were transferred from half-strength MS agar solidified medium (1.5% sucrose) plates to half-strength MS agar solidified medium (1.5% sucrose) for control plates and plates supplemented with 50 and 100 mM of NaCl. Seedlings were transferred to NaCl plates after 2 days (etiolated seedlings) or 3 days (light-grown seedlings) and photographed 3 days or 7-8 days later, respectively.

#### Plasmid constructs

A genomic fragment spanning the 1.4-kb *TTL1* promoter region upstream of the start codon and the *TTL1* genomic region without stop codon was PCR amplified using the primers detailed in Supplementary Table S1 and cloned into the pENTR/D-TOPO vector (Invitrogen). The CESA1-C coding sequence for expression in *N. benthamiana* was PCR amplified using specific primers using pGBKT7-CesA1-C domain as template and the attB1 adapter universal primer (Invitrogen) and introduced into the pDONR/Zeo vector (vector) using the BP Clonase II Gateway cloning kit (Invitrogen). The truncated version TTL3IDR was generated using the Q5 Site-Directed Mutagenesis Kit (New England Biolabs) to remove base pairs 610 to 2071 of the TTL3 CDS, using pENTR/D-TOPO-TTL3 full length CDS as template. All the resulting pENTR clones were verified by diagnostic PCR, restriction analysis, and sequencing. These pENTR clones in combination with the appropriate destination vectors (pDEST) were used to create the final Gateway-expression constructs by Gateway LR Clonase II reaction (Invitrogen).

The pGWB4, pGWB5 and pGWB14 vectors, from the pGWB vector series, were provided by Tsuyoshi Nakagawa (Department of Molecular and Functional Genomics, Shimane University; Nakagawa et al., 2007) and were used as pDEST for either transient expression in *Nicotiana benthamiana* or generating stable lines in *A. thaliana*. Finally, the Y2H pGADT7(GW) and pGBKT7(GW) destination vectors were provided by Salomé Prat (Centro Nacional de Biotecnología-Consejo Superior de Investigaciones Científicas).

The 35S::TTL3-GFP, 35S::BAK1-HA, as well as the TTL3, TTL3Δ4TPR and TTL3ΔIDR Y2H constructs were described previously (*4*) and used in this study. The CESAs constructs used for the Y2H assay were generated by amplifying the N-terminal and internal catalytic loop of *CESA1 (*CESA1-N:1-270aa; CESA1-C: 320-856aa), *CESA3* (CESA3-N: 1-260aa; CESA3-C: 304-842aa) and *CESA6 (*CESA6-N: 1-277aa; CESA6-C: 321-868aa), identified based on transmembrane topology predictions, from *Arabidopsis thaliana* Col-0 cDNA with primers listed in Table S1. The fragments and the matchmaker Gold Y2H vector pGBKT7 were digested (NdeI/SfiI), ligated, and transformed into Saccharomyces cerevisiae strain AH109 as described in the “Yeast Two-Hybrid Assay” section.

Plasmids for heterlogous protein expression of full length TTL3 and 6xHis-TTL3IDR were constructed as follows. The vector petM11SUMO3GFP was cut with BamHI and the full length TTL3 CDS, amplified with specific overhangs (Table S1) was inserted using the Gibson assembly method (*8*). The TTL3 IDR (aa 1-200) was amplified with specific overhangs (Table S1) for the pPROEX HTb vector (Invitrogen, USA) and fused with the linearized vector (EcoRI + XbaI) using the Gibson assembly technique.

#### Generation of transgenic plants

Expression clones were transformed into *Agrobacterium tumefaciens* strain GVG3101::pMP90. The pGWB4 harbouring the full length line *pTTL1::TTL1g-GFP* construct was transformed into *ttl1ttl3 Arabidopsis* plants by floral dipping (*9*) to generate stable transgenic plants. Seeds obtained were sowed in half-strength MS agar-solidified medium supplemented with 1.5% sucrose [w/v] and with 50 μg/ml Hygromycin B (Duchefa). T4 homozygous transgenic plants were used in this study.

#### Transient expression in *N. benthamiana*

Transient expression in *N. benthamiana* was performed following the protocol described by (*10*). A*. tumefaciens* (GV3101::pMP90) carrying the different constructs was used together with the p19 strain. Cultures were grown overnight in Luria-Bertani (LB) medium containing rifampicin (50 mg/ml), gentamycin (25 mg/ml) and antibiotic specific for the positive selection of the plasmid constructs. Individual colonies were used for inoculation on LB for a given construct. After overnight incubation at 28°C, *Agrobacterium* cells were collected by centrifugation (15 min at 3000g in 50-ml falcon tubes) at room temperature and pellets were resuspended in agroinfiltration solution (10 mM MES, pH 5.6, 10 mM MgCl2 and supplemented with 100 μM acetosyringone). Subsequently, cultures were incubated for 2-4 h at room temperature in dark conditions. Bacterial suspension was adjusted at OD_600_ of 0.8 for the constructs and 0.2 for the p19 strain to reach a final OD_600_ of approximately 1 for agroinfiltration. For double infiltration experiments, Agrobacterium strains were infiltrated at OD_600_ of 0.4 for each construct and at OD_600_ of 0.2 for the p19 strain. Cultures were infiltrated in 4-week-old *N. benthamiana* leaves using 1ml syringes (BD Plastipak) at the abaxial side of the leaf. After infiltration, all plants were kept in the greenhouse and analysed 2 days later.

#### Protein extraction and Co-IP in *N. benthamiana*

Protein extraction and Co-IP in *N. benthamiana* were performed as described previously (*11*) with some modifications. Briefly, 4-week-old *N. benthamiana* plants were used for transient expression assays as described in “Transient Expression in *N. benthamiana”* section. *N. benthamiana* leaves were mainly ground to fine powder in liquid nitrogen. Approximated 0.5 g of ground leaves per sample was used, and total proteins were then extracted with extraction buffer (50 mM Tris-HCl, pH 7.5, 150 mM NaCl, 10% glycerol, 10 mM EDTA pH8, 1 mM NaF, 1 mM Na_2_MoO_4_·2H_2_O, 10 mM DTT, 0.5 mM PMSF, 1% [v/v] P9599 protease inhibitor cocktail [Sigma-Aldrich]); Nonidet P-40, CAS: 9036-19-5 [USB Amersham Life Science] 0.5% (v/v), added at 2 ml/g powder using an end-over-end rocker for 30 min at 4°C. Samples were centrifuged 20 min at 4°C and 9000 rpm (9056 g). Supernatants (approx. 4 mg/ml protein) were filtered by gravity through Poly-Prep chromatography columns (731-1550, Bio-Rad), and 100 μL was used as input. The remaining supernatants were incubated for 2 h at 4°C with 15 μL of GFP-Trap coupled to agarose beads (Chromotek) in an end-over-end rocker. During incubation of protein samples with GFP-Trap beads, the final concentration of detergent (Nonidet P-40) was adjusted to 0.2% (v/v) in order to avoid nonspecific binding to the matrix as recommended by the manufacturer. Subsequent incubation, beads were collected and washed four times with the wash buffer (similar to extraction buffer but without detergent). Finally, beads were resuspended in 75 ml of 2X concentrated Laemmli sample buffer and heated at 70°C for 20 min to dissociate immunocomplexes from the beads. Total (input), immunoprecipitated (IP), and CoIP proteins were separated in a 10% SDS-PAGE gel and analysed as described in the section “Immunoblot”.

#### Spinning disk live cell imaging and data processing

3-day-old plant hypocotyls or 5-day-old roots were covered with a 1% agarose cushion and mounted onto the imaging system as described previously (*12*). XFP(Fluorescence?)-tagged proteins were imaged with a CSU-W1 T1 Yokogawa spinning disk head fitted to a Nikon Eclipse Ti2-E-inverted microscope with a CFI Apo TIRF × 100 N.A. 1.49 oil immersion objective and two iXon Life 888 EM-CCD cameras (Andor, GB). The system was assembled by 3i (https://www.intelligent-imaging.com). GFP was imaged using a 488 nm solid-state diode laser (150 mW) and a 525/30-25 nm emission filter and RFP was detected with a 561 nm solid-state diode laser (150 mW) and a 617/73-25 nm emission filter. Alternatively, a CSU-W1 Yokogawa spinning disk head was fitted to a Nikon Eclipse Ti-E-inverted microscope with a CFI PlanApo × 100 N.A. 1.40 oil immersion objective and two iXon Ultra EM-CCD cameras (Andor, GB). For this visitron system (https://www.visitron.de), GFP was imaged using a 488 nm solid-state diode laser (200 mW) and a 525/50 nm emission filter; RFP was detected with a 561 nm solid-state diode laser (200 mW) and a 630/75 nm emission filter. Time lapse images were processed and analyzed with Fiji (*13*). Drifts were corrected by using the plugin StackReg or MultiStackReg in cases where two channels were imaged (*14*). When the drift of samples could not be corrected in this way, they were excluded from the analysis. Backgrounds were subtracted by the “Subtract Background” tool (rolling ball radius, 30–50 pixels). To quantify CesA velocities, three frames were averaged by “WalkingAverage” and kymograph analysis was performed with the kymograph tool of FIESTA (*15*).

#### Microtubule quantification, particle detection and particle clustering at the plasma membrane

Particle density at the plasma membrane of cells was determined as described before (*16, 17*). Microtubule density measurements were done as described in detail before (*18*). Particle clustering at the plasma membrane (e.g. by recruitment of cytosolic TTL3-GFP to the plasma membrane localized CESA particles) was assessed by measuring the skewness of the image histogram with Fiji. A predefined cell area was cut-out from all analyzed cells. The area was designed to fit into all analyzed cells and thereby excluding all non-cell signals (e.g. the area outside of the analyzed cell and the outer cell lines). All images were taken with constant settings and raw images were used for this analysis to not affect the histogram.

#### Yeast Two-Hybrid Assay

The Gal4-based yeast two-hybrid system (Clontech Laboratories) was used for testing the interaction of TTL3Δ4TPR, TTL3ΔIDR and TTL3IDR with the N-terminal domains (CESA-N) and the cytosolic catalytic domains (CESA-C) of CESA1, CESA3, and CESA6. The bait and prey constructs are detailed in the “Plasmid Constructs” section. Bait and prey plasmids were transformed into Saccharomyces cerevisiae strain AH109 with the lithium acetate/single-stranded carrier DNA/polyethylene glycol method and transformants were grown on plasmid-selective media (synthetic defined (SD)/–Trp–Leu). Plates were incubated at 28°C for 4 days and independent colonies for each bait–prey combination were resuspended in 200 μL of sterile water. Ten-fold serial dilutions were made and 5 μL of each dilution were spotted onto three alternative interaction-selective medium (SD/–Trp–Leu–His+3-AT (for 3-amino-1,2,4-triazole, 2mM), SD/–Trp–Leu-Ade, and SD/–Trp–Leu–Ade+3-AT). Plates were incubated at 28°C and photographed 3 or 7 days later.

#### Yeast Two-Hybrid Protein Extraction

For immunoblot analysis, one or two independent yeast co-transformants (a and b) for each bait–prey plasmid combination were grown in 50 ml of SD/–Leu–Trp to an OD600 of 0.7 to 1. Cultures were centrifuged at 4000 rpm for 3 min. The resulting pellet was washed once with cold water and re-suspended in 200 ml of RIPA buffer (2 mM sodium phosphate buffer, pH 7, 0.2% Triton X-100, 0.02% [w/v] SDS, 0.2 mM EDTA, pH 8, and 10 mM NaCl) containing protease inhibitor (1 tablet/10 ml, cOmplete, Mini, EDTA-free Protease Inhibitor Cocktail, Roche). Glass beads (500 ml, 425 to 600 mm, Sigma-Aldrich) were added, and the sample was vortexed in FastPrepTM FP120 (BIO 101) at a power setting of 5.5 for two 15-s intervals separated by 1-min interval on ice. Next, 400 ml of RIPA buffer with protease inhibitors was added, and the sample was vigorously vortexed. The supernatant was recovered, and the protein concentration was determined using Bradford assays. Total protein (50 μg) was resolved on 10% polyacrylamide/SDS gels and analyzed by immunoblotting to detect the Gal4AD-HA fusion proteins with an anti-HA antibody (1:5000, Sigma, H3663) as described in the “Immunoblot’’ section.

#### Preparation of total, soluble and microsomal fractions for immunoblot

Three-day-old etiolated *ttl1ttl3ttl4 TTL3-GFP* (Fig. S1K), *ttl1ttl3ttl4 TTL3-GFP* and *prc1-1 TTL3-GFP* (Fig. 3E) seedlings grown in half-strength solidified MS agar with 1% (w/v) sucrose, were used for subcellular fractionation analysis. For the subcellular fractionation analysis under salt stress (Fig3B), *ttl1ttl3ttl4 TTL3-GFP* etiolated seedlings were grown for 3 days in half-strength MS agar solidified medium with 1% (w/v) sucrose and then transferred to half-strength MS liquid medium with 1% (w/v) sucrose (control) or supplemented with 200 mM NaCl. Samples were collected 0.5 and 28 hours after the salt stress imposition.

The total, microsomal and soluble fractions were prepared using a modified method that allows the use of a limited amount of tissue sample (*19*) with some modifications. Briefly, 100 mg of 3-day-old etiolated seedlings were grounded in 1.5 ml microcentrifuge tubes using polypropylene pestles with 1.5x EB buffer on ice. The concentrated 1.5x EB was used to account for tissue water content of etiolated seedlings. The 1.0x EB consisted of 100 mM Tris– HCl (pH 7.5), 25% (w/w, 0.81 M) sucrose, 5% (v/v) glycerol, 10 mM ethylenediaminetetraacetic acid (EDTA, pH 8.0), 5 mM ethyleneglycoltetraacetic acid (EGTA pH 8.0), 5 mM KCl, and 1 mM 1,4-dithiothreitol (DTT). To inhibit proteolysis and phosphatase activity the 1.5x EB was supplemented with 0.2% (w/v) bovine serum albumin (BSA), 50 mM sodium fluoride (NaF), 2 mM sodium molybdate (Na2MoO4), 1 mM Pefabloc-SC (AEBSF) and 1% (v/v) protease inhibitor cocktail (ref: P9599, Sigma-Aldrich). The homogenate was transferred to the prepared PVPP pellets that were previously equilibrated in 200 mM Tris– HCl (pH 7.5), 40% (w/w, 1.37 M) sucrose, 20 mM EDTA (pH 8.0), 20 mM EGTA (pH 8.0), 10 mM KCl, and 0.4% BSA, mixed (vortex 30 sec), and left for 5 min for the PVPP to adsorb phenolic compounds. Samples were cleared by centrifugation (600g, 3 min 4°C). The supernatants were saved, and pellets were re-extracted twice using 1.1x EB and 1.0 EB, respectively. Combined supernatants were re-cleared (600g, 3 min at 4°C), aliquots for total protein were saved and combined supernatants were centrifuged (21000g for 1h 40min at 4°C) to yield total membrane pellets. Supernatants were saved for the soluble protein fraction analysis. The membrane pellets were washed with wash buffer that consist of 20 mM Tris–HCl (pH 7.5), 5 mM EDTA, 5 mM EGTA, 1 mM PMSF and 1% (v/v) protease inhibitor cocktail and centrifuged (21000g, 45 min). Then pellets were resuspended in storage buffer that consist of 20 mM Tris–HCl pH 7.5, 5 mM EDTA, 5 mM EGTA, 20% glycerol, 1mM DTT, 50 mM NaF, 2 mM Na2MoO4, 1 mM AEBSF and 1% (v/v) protease inhibitor cocktail. For immunoblot analysis, total and soluble fractions were 1x concentrated compared with microsomal protein extract. Samples were diluted in 2x Laemmli buffer (125 mM Tris-HCl, pH 6.8, 4% [w/v] SDS, 20% [v/v] glycerol, 2% [v/v] b-mercaptoethanol, and 0.01% [w/v] bromophenol blue) denatured by heating samples for 45 min at 70°C, separated in a 10% SDS-PAGE gel and analyzed as described in the section “Immunoblot”.

#### Immunoblot analysis

Proteins separated by SDS–PAGE polyacrylamide gel electrophoresis were blotted using Trans-blot Turbo Transfer System (Bio-Rad) or wet electroblotting systems (Bio-Rad) onto polyvinylidene difluoride (PVDF) membranes (Immobilon-P, Millipore) following instructions by the manufacturer. PVDF membranes, containing electroblotted proteins, were then incubated with the appropriate primary antibody followed by the appropriate secondary peroxidase-conjugated antibody. The following primary antibodies were used for detection of epitope-tagged proteins: mouse monoclonal anti-GFP clone B-2 (1:1000, catalog no. sc-9996, Santa Cruz Biotechnology) mouse monoclonal anti-HA clone HA-7 (1:3000, catalog no. H3663, Sigma-Aldrich) and rabbit polyclonal anti-SYT1 antibody (1:5000). The secondary antibodies used in this study were anti-mouse IgG whole molecule-Peroxidase (1:80000; catalog no. A9044, Sigma-Aldrich) and anti-rabbit IgG whole molecule-Peroxidase (1:80000; catalog no. A0545, Sigma-Aldrich). Proteins and epitope-tagged proteins on immunoblots were detected using the Clarity ECL Western Blotting Substrate or SuperSignal West Femto Maximum Sensitivity Substrate according to the manufacturer’s instructions, and images of different time exposures were acquired using the Chemidoc XRS1 System (Bio-Rad). Only images with no saturated pixels were used for protein quantification. Immunoblotted PVDF membranes were stained with Coomassie Brilliant Blue R-250 to confirm equal loading of the different samples in each experiment.

#### Heterologous protein expression

6xHis-TTL3IDR and 6xHis-SUMO3-TTL3 were expressed in ArcticExpress (DE3) E. coli Cells (Agilent). A starter culture was grown overnight at 28 °C and used to inoculate the main cultures in a ratio of 1:10. Cultures were grown at 37 °C until an OD600 ~0.6 was reached. Cultures were moved to 12 °C and protein expression was induced when cultures were cooled down by addition of isopropyl β-D-1-thiogalactopyranoside (IPTG) at a final concentration of 1 mM. Cells were collected after overnight incubation by centrifugation at 5,000 *xg* and washed in 150 mM NaCl. Pellets were resuspended in lysis buffer (50 mM Tris-HCl, 200 mM NaCl, 20 mM imidazole, pH 7.4 for 6xHis-TTL3IDR and 50 mM Hepes, 200 mM NaCl, 20 mM imidazole, pH 7.2 for 6xHis-SUMO3-TTL3). From here on, all following Tris based buffers were used for 6xHis-TTL3IDR and Hepes based buffers for 6xHis-SUMO3-TTL3. Cell lysates were prepared by passing the solution through an Microfluidics M-110P Homogenizer (Microfluidics Corp., USA) at 22,000 psi for four times. Cellular debris was spun down at 20,000 *xg*, supernatant was collected and filtered through a 0.2 μm syringe filter. Recombinant proteins were purified using Ni Sepharose™ High Performance HisTrap™ HP columns (GE Healthcare Life Sciences, USA) and an ÄKTA pure 25L (GE Healthcare Life Sciences, USA) equipped with a sample pump. Buffers for protein purification were prepared as follows: Buffer A: 50 mM Tris-HCl (pH 7.4), 200 mM NaCl or 50 mM Hepes (pH 7.2), 200 mM NaCl; Buffer B: 50 mM Tris-HCl (pH 7.4), 200 mM NaCl, 500 mM Imidazole or 50 mM Hepes (pH 7.2), 200 mM NaCl, 500 mM Imidazole. Sample application onto the column was performed at 1 ml/min with 96% A and 4% B. The column was washed stepwise at 1 ml/min as follows: 20 column volumes (CV) 96% A and 4% B, 20 CV 90% A and 10% B. Final protein sample was eluted at 1 ml/min for 20 CV with a linear gradient starting at 90% A and 10% B until 0%A and 100% B were reached. Elution was performed in upflow mode. Samples were collected in 2 ml fractions. Fractions enriched with protein were combined and gel filtered to remove imidazole using PD-10 Desalting Columns (GE Healthcare, USA) according to the gravity protocol in the manual. The column was equilibrated with 50 mM Tris-HCl (pH 7.4), 200 mM NaCl or 50 mM Hepes (pH 7.2), 200 mM NaCl. Proteins were concentrated at 12 °C and 3,000 *xg* using Macrosep® Advance Centrifugal Devices with a 3K or 30K cutoff (Pall Corporation, USA). 6xHis-TTL3IDR was snap frozen and stored at −80 °C until further use. 6xHis-SUMO3-TTL3 was cut with SUMO protease (bdSENP1 (*20*)) in a ratio of 500:1 at 12 °C for 4h under gentle agitation. The cleavage products were separated by size exclusion chromatography using a HiLoad 16/600 Superdex 200 pg (Cytiva, U.S.A.) column. The column was equilibrated in storage buffer (50 mM Hepes (pH 7.2), 200 mM NaCl, 10% glycerol) and cleavage products were separated at a flow rate of 1 ml/min for 1.5 column volumes. Fraction collection was initiated after 0.2 column volumes. The final protein was concentrated as described above and snap frozen until further use.

#### Microtubule affinity and turbidity assays

Microtubule spin down and turbidity assays were performed as described in detail before (*16, 21*). Briefly, to determine the dissociation constant, a constant amount (7.5 μg) of 6xHis-TTL3IDR or TTL3 was incubated with increasing concentrations of polymerized microtubules ranging from 0 to 7.2 μM (calculated based on the MW of a tubulin dimer). 0.05% Tween 20 was added to all reactions. The samples were incubated for 30 min at RT and subsequently spun at 25,000 × g for 30 min at RT to pellet microtubules and bound proteins. Supernatant and pellet fractions were subjugated to SDS-PAGE and protein levels in both supernatant and pellet fractions were analyzed using the Gel-function of Fiji. Final dissociation constant (KD) was estimated by fitting a saturation binding curve onto the data points with GraphPad Prism v9.0 (GraphPad software, Inc., USA).

#### Phylogenetic analysis

Marchantia polymorpha and Physcomitrella patens were used as representatives from embryophytes (earliest land plants). Arabidopsis thaliana TTL3 protein sequence was used as input for a PSI-BLAST analysis against Marchantia polymorpha, Physcomitrella patens, and all the NCBI organisms tagged as chlorophyta (taxid:3041) and charophytes (taxid:3146). In the case of Marchantia polymorpha and Physcomitrella patens results with the smallest E-value with IDR, TPR and TRXL (a region with homology to class h thioredoxins but lacking essential Cys residues required for thioredoxin activity) domains present in the same sequence, were used as representatives. For chlorophyta and charophytes no results were obtained with IDR, TPR and TRXL domains. Thus, in the case of chlorophyta, the 4 results with the smallest Evalue and with TPR and TRX (a region with homology to class h thioredoxins and conserving the essential Cys residues required for thioredoxin activity) domains were used as representatives, meanwhile for charophytes the only results came from Chara braunii. Domains were retrieved from InterPro, using the outputs from MOBIdb_Lite (IDR), SMART (TPR), CDD (TRX) and InterPro (Chara braunii domains). Domains were illustrated using IBS 1.0.3 (*22*)and Adobe Illustrator CC 2017 21.0.0. Alignment was carried out using the web versión of (*23, 24*) MAFFT version 7 (https://mafft.cbrc.jp/alignment/server/) with standard parameters, and the phylogenetic analysis was performed with desktop MEGA-X tool (*25*) (ver. 10.1.7, Neighbor-Joining method, Bootstrap method, 500 replications [numbers from 0 to 100, substitution model: p-distance [numbers from 0 to 1], uniform rate among sites, gaps treatment: pairwise deletion).

#### Statistical analysis and experimental design

For statistical analyses, Welch’s unpaired t-test, two-tailed unpaired t-test, one-way ANOVA, Welch’s ANOVA or two-way ANOVA were performed using GraphPad Prism 9.3. If appropriate, Dunnett’s T3 multiple comparisons test or Tukey’s multiple comparisons test were performed afterwards. A *p*-value of < 0.05 was considered as statistically significant. Statistical methods used to calculate *p*-values are described in the figure legends. Data analysis (especially for images) was either done automatically or file names were removed before the analysis. For the measurement of plant size, investigators were not blinded. In this case, data were always collected according to the genotype of plants. Sample size was determined for each experiment based on similar data reported in scientific literature.

---

**Fig S1.**
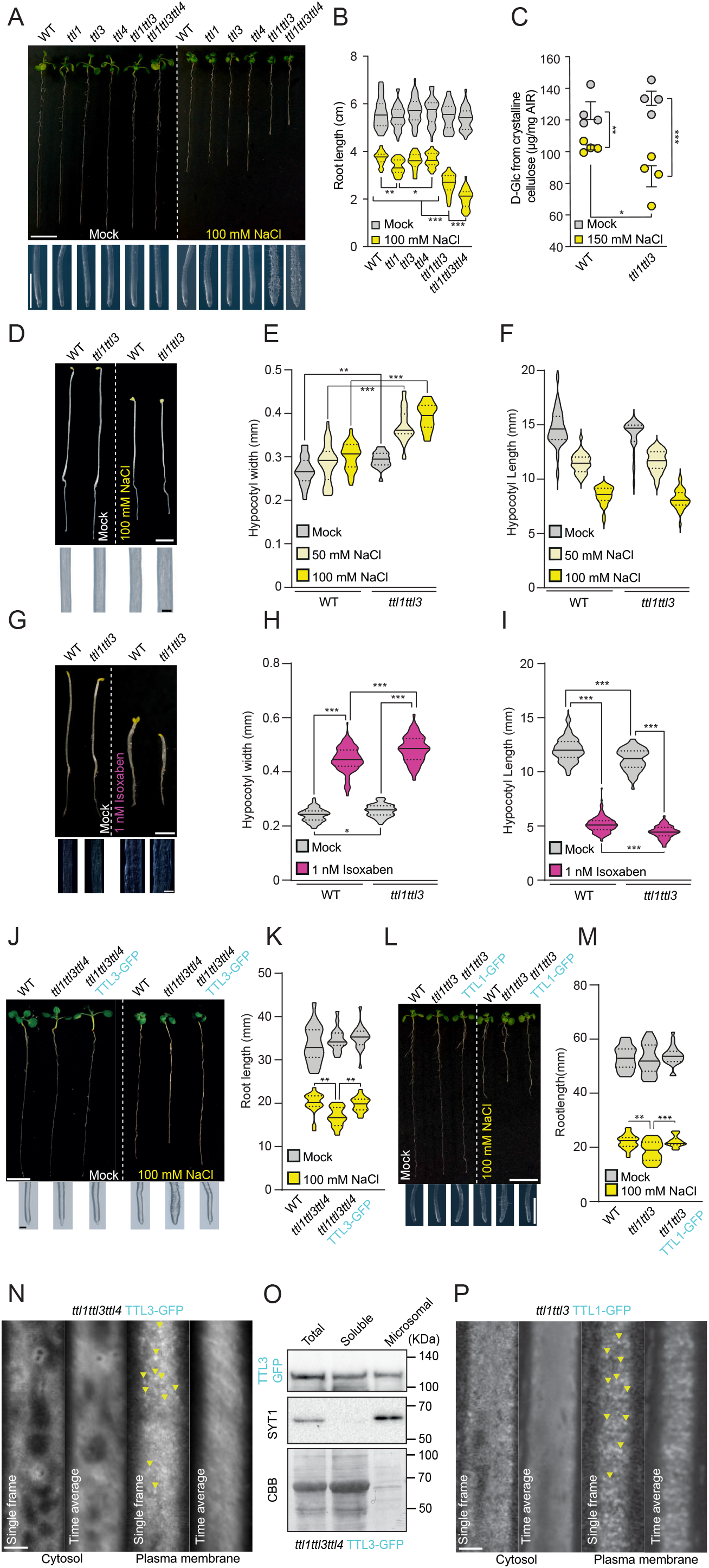
TTL1 and TTL3 are peripheral membrane proteins required for plant tolerance to salt stress and are associated with cellulose synthesis. (**A**) Wild-Type (Col-0) and *ttl* mutants were germinated and grown for 3 days in control conditions and then transferred to control media or media supplemented with 100 mM NaCl and grown for additional 8 days. Magnifications show distinctive root tip swelling in *ttl* double and triple mutant plants under salt stress, indicative of defective anisotropic growth (lower panels). Scale bars=1 cm (upper panels) and 1 mm (lower panels). (**B**) Root length quantification of seedlings as in (A). Violin plots: centerlines show the medians; dotted lines indicate the 25th and 75th percentiles; N≥27 roots and three independent experiments. Welch’s ANOVA between 100 mM NaCl treated groups: *p*-value≤0.001; Dunnett’s T3 multiple comparison: \**p*-value≤0.05, \*\**p*-value≤0.01, \*\*\**p*-value≤0.001. (**C)** Cellulose represented as μg of D-glucose derived from cellulose per mg of dried alcohol-insoluble residue (AIR) of roots grown as depicted in (A) but using 150 mM NaCl. Values are mean±SEM; N=4 biological replicates; 2 technical replicates per biological replicate. Twotailed, unpaired t-test; \**p*-value≤0.05, \*\**p*-value≤0.01, \*\*\**p*-value≤0.001. (**D**) Dark grown WT and *ttl1ttl3* seedlings germinated for 2 days in control conditions and transferred to either control plates or plates supplemented with 100 mM NaCl and grown for additional 3 days in the dark. Magnifications show distinctive hypocotyl width in *ttl1ttl3* double mutant plants under salt stress (lower panels). Scale bar=2 mm (upper image) and 200 μm (close-up hypocotyls). (**E**) and (**F**) Quantification of hypocotyl width and length of seedlings as depicted in (D). Violin plots: centerlines show the medians; dotted lines indicate the 25th and 75th percentiles; N≥18 seedlings from 3 independent experiments. Welch’s unpaired t-test; \**p*-value≤0.05, *\*\*p-* value≤0.01, \*\*\**p*-value≤0.001. (**G**) Dark grown WT and *ttl1ttl3* seedlings germinated on control or media supplemented with 1 nM isoxaben for 5 days in the dark. Magnifications show distinctive hypocotyl width in *ttl1ttl3* double mutant plants in 1nM isoxaben, indicative of defective anisotropic growth (lower panels). Scale bar=5 mm (upper image) and 200 μm (close-up hypocotyls). (**H**) and (**I**) Quantification of hypocotyl width and length of seedlings as depicted in (G). Violin plots: centerlines show the medians; dotted lines indicate the 25th and 75th percentiles; N≥11 seedlings from 3 independent experiments. Unpaired t-test; \**p*-value≤0.05, \*\**p*-value≤0.01, \*\*\**p*-value≤0.001. (**J**) The *TTL3-GFP* construct complements the root growth defects of the *ttl1tt3ttl4* mutant. seedlings on NaCl. WT (Col-0), *ttl1ttl3ttl4*, and *ttl1ttl3ttl4* TTL3-GFP seedlings were germinated and grown on control conditions and transferred after 3 days to control and NaCl containing media and grown for an additional 8 days in the light. Magnifications (lower panel) show distinctive root tip swelling in *ttl1ttl3ttl4* triple mutant plants under salt stress, indicative of anisotropic growth failures. Scale bars=5 mm (upper image) and 200 μm (close-up root tips). (**K**) Quantification of root length of seedlings as in (J). Violin plots: centerlines show the medians; dotted lines indicate the 25th and 75th percentiles; Welch’s ANOVA between mock groups: *p*-value=0.29; Welch’s ANOVA between NaCl treated groups: *p*-value≤0.001; Dunnett’s T3 multiple comparison: \*\**p*-value≤0.01. N≥15 roots and three independent experiments. (**L**) The *TTL1-GFP* construct complements the root growth defects of the *ttl1tt3* mutant. seedlings on NaCl. WT (Col-0), *ttl1ttl3*, and *ttl1ttl3* TTL1-GFP seedlings were germinated and grown on control conditions and transferred after 3 days to control and NaCl containing media and grown for an additional 8 days in the light. Magnifications (lower panel) show distinctive root tip swelling in *ttl1ttl3* triple mutant plants under salt stress, indicative of anisotropic growth failures. Scale bars=1 cm (upper image) and 1 mm (close-up root tips) μm. (**M**) Quantification of root length of seedlings as in (L). Violin plots: centerlines show the medians; dotted lines indicate the 25th and 75th percentiles; Welch’s ANOVA between mock groups: *p*-value=0.29; Welch’s ANOVA between NaCl treated groups: *p*-value≤0.001; Dunnett’s T3 multiple comparison: \*\**p*-value≤0.01, \*\*\**p*-value≤0.001. N≥15 roots and two independent experiments. (**N**) TTL3-GFP in 3-day-old hypocotyl epidermal cells of *ttl1ttl3ttl4* TTL3-GFP seedlings. TTL3-GFP signal is visible as cytosolic signal and as distinctive motile particles (yellow arrows) at the plasma membrane as revealed by single frames and time-average projections. Scale bar=2.5 μm. (**O**) Total, soluble, and microsomal fractions were isolated from 3-day-old *ttl1ttl3ttl4* TTL3-GFP etiolated seedlings. A specific antibody for the microsomal SYT1 protein as a control of the purified fraction was used to detect this microsomal control protein. Coomassie brilliant blue (CBB) was used to confirm equal loading. The experiment was repeated 3 times with similar results. (**P**) TTL1-GFP in 3-day-old root epidermal cells of *ttl1ttl3* TTL1-GFP seedlings. TTL1-GFP signal is visible as cytosolic signal and as distinctive motile particles (yellow arrows) at the plasma membrane as revealed by single frames and time-average projections. Scale bar=2.5 μm.

**Fig. S2.**
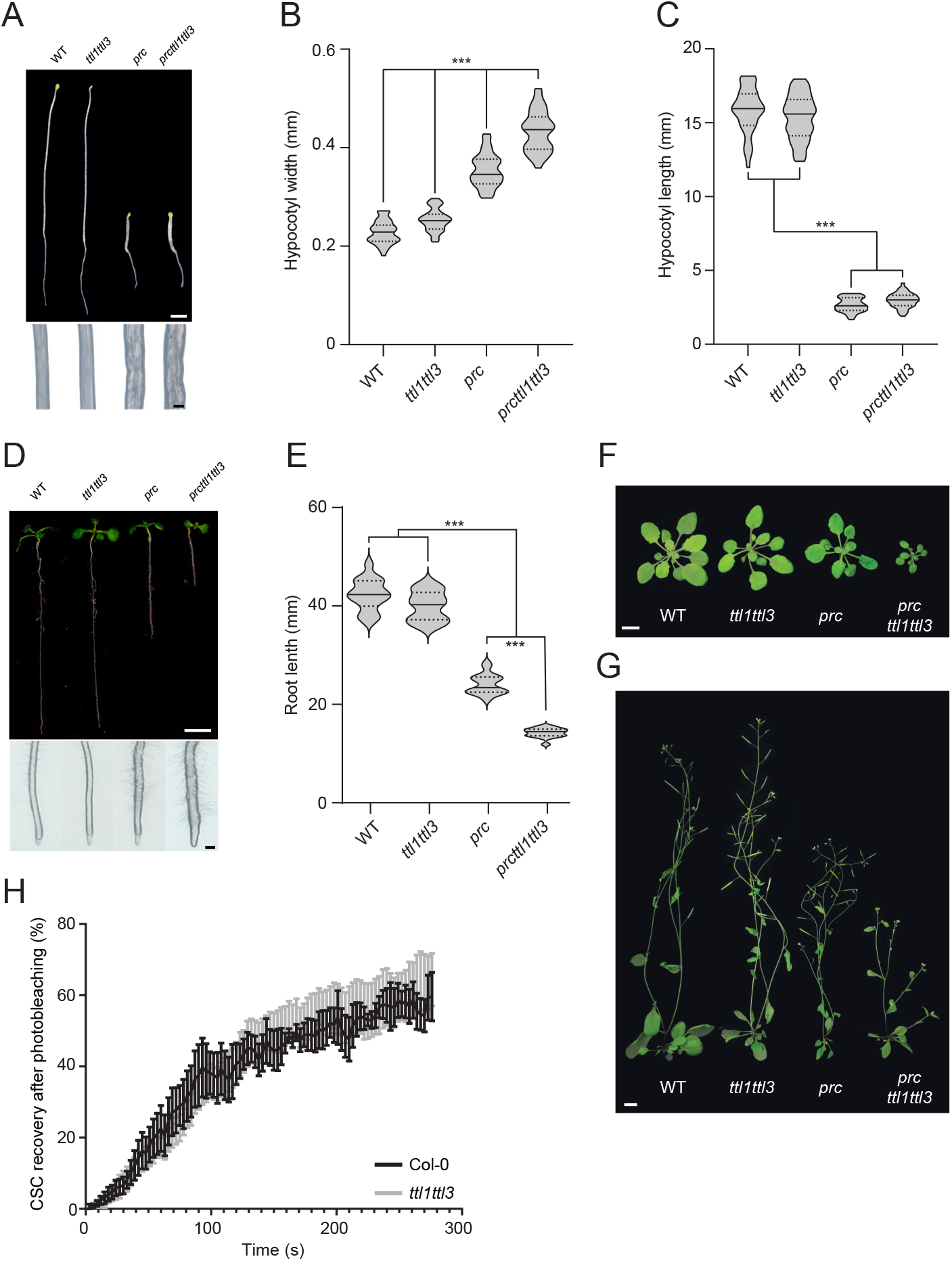
*ttl1ttl3* exacerbates the *prc1-1* developmental defects and does not affect CESA delivery to the plasma membrane. (**A**) Dark grown WT (Col-0), *ttl1ttl3, prc1-1*, and *ttl1ttl3prc1-1* seedlings grown in control conditions for 5 days. Magnifications show distinctive hypocotyl width in *prc1-1* and *prc1-1ttl1ttl3* triple mutant plants when compared to WT (Col-0) and *ttl1ttl3*, indicative of defective anisotropic growth (lower panels). Scale bar=2mm (upper image) and 200 μm (close-up hypocotyls). (**B**) and (**C**) Hypocotyl width and length quantification of seedlings as depicted in (A). Violin plots: centerlines show the medians; dotted lines indicate the 25th and 75th percentiles; Welch’s ANOVA on hypocotyl width: *p*-value≤0.001; Welch’s ANOVA on hypocotyl length: *p*-value≤0.001; Dunnett’s T3 multiple comparisons: \*\*\**p*-value≤0.001; N≥35 seedlings and two independent experiments. (**D**) Ten-day-old WT (Col-0) and *ttl1ttl3, prc1-1*, and *ttl1ttl3prc1-1* seedlings grown on control media (upper panels) in the light. Magnifications (lower panel) show distinctive root tip swelling in *prc1-1* and *ttl1ttl3prc1-1* triple mutant plants indicative of isotropic growth defects. Scale bars=1 cm (upper image) and 250 μm (close-up root tips). (**E**) Quantification of root length of seedlings in (D). Violin plots: centerlines show the medians; dotted lines indicate the 25th and 75th percentiles; Welch’s ANOVA on hypocotyl length: *p*-value≤0.001; Dunnett’s T3 multiple comparisons: \*\*\**p*-value≤0.001; N≥16 roots and three independent experiments. (**F**) and (**G**) Morphological phenotypes of 5-week-old plants of the indicated genotypes grown in either short-day (F) or long-day conditions (G). Scale bar=1 cm. (**H**) Fluorescence Recovery After Photobleaching (FRAP) of 3-day-old WT and *ttl1ttl3* etiolated hypocotyl epidermal cells expressing YFP-CESA6. Graph displays re-population of the plasma membrane at indicated time points with fluorescent CESA foci as percentage of CESA density at t=0 (just before FRAP). Two-way ANOVA analysis of CESA recovery; *p*=0.84 (genotype), *p*≤0.001 (time), *p*>0.99 (genotype × time). n≥7 cells from at least five seedlings and three independent experiments. Values are mean±SEM.

**Fig. S3.**
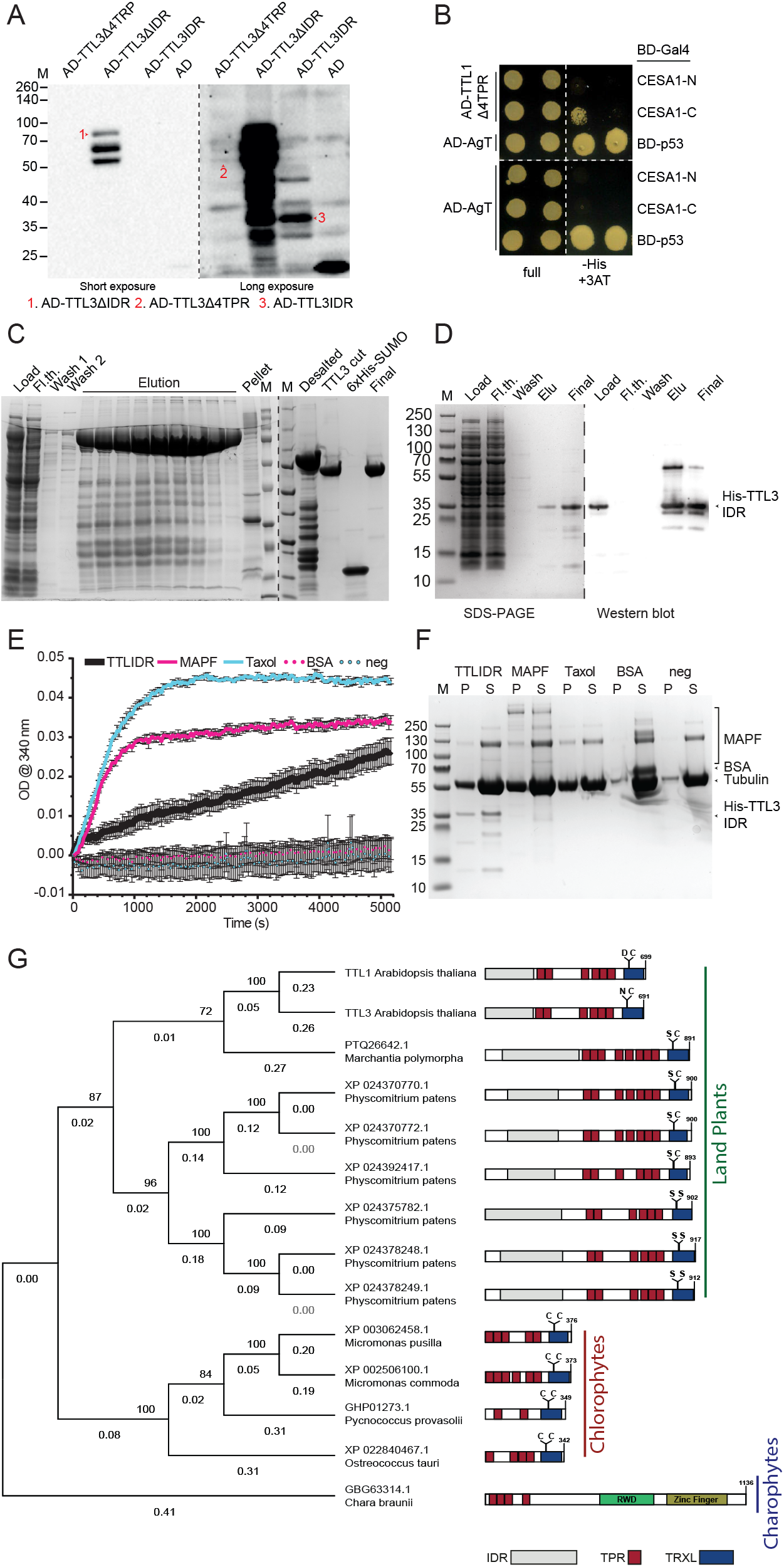
The TTL1 N-terminus interacts with the catalytic domain of CESA1 and the TTL3 IDR interacts with and promotes microtubule formation. TTL proteins have emerged in land plants. (**A**) Immunoblot analysis showing the expression of AD-TTL3ΔIDR, AD-TTL3Δ4TPR, and AD-TTL3IDR in protein extracts of yeast transformants used for the Yeast-two-hybrid assays (shown in Fig. 3B). Yeast transformants for each were resolved in polyacrylamide/SDS-Page gels and analyzed by immunoblot using an anti-HA Tag monoclonal antibody. HA tag is transcriptionally fused to AD (Activation Domains). The expected molecular size of (1) AD-TTL3ΔIDR, (2) AD-TTL3Δ4TPR and (3) AD-TTL3IDR is represented in the figure. (**B**) Yeast-two-hybrid assays showing the interaction of TTL1Δ4TPR (amino acids 1 to 307) with CESA1-C. Growth on non-selective media (full) and interaction-selective media (lacking histidine [–His] and supplemented with 3-amino-1,2,4-triazole [+3AT]) are shown. Photographs were taken after 3 days. A positive control (BD-p53/AD-AgT) for the yeast two-hybrid assays is shown. -N=N-terminal region of CESAs; -C=catalytic cytosolic loop of CESAs. The experiment was repeated 3 times with the same results. (**C**) Expression and purification of 6xHis-SUMO-TTL3. SDS-PAGE gel of the purification steps. M = Molecular marker; Load = crude protein extract; Fl.th. = flowthrough after Ni-NTA Agarose binding; TTL3 cut = TTL3 after digestion with SUMO protease and gel filtration; Final = Desalted, cut and concentrated protein. (**D**) Expression and purification of 6xHis-TTL3IDR. Left panel: Coomassie-stained SDS-PAGE gel of the purification steps. M = Molecular marker; Load = crude protein extract; Fl.th. = flowthrough after Ni-NTA Agarose binding; Elu = Imidazole elution; Final = Desalted and concentrated protein. Right panel: Representative image of a western blot to detect 6xHis-TTL3IDR. **(E)** Microtubule turbidity assay. Tubulin was incubated with buffer or BSA as negative controls, a microtubule associated protein fraction (MAPF) or taxol as positive controls, and 6xHis-TTL3IDR. Microtubule formation was measured at 340 nm. Values are mean±SEM. N=3 technical replicates. **(F)** Reactions, as shown in (E), were spun down, separated in pellet and supernatant fractions, and analyzed by SDS-PAGE. While distinct tubulin bands at 55 kDa are visible in the pellet fraction of the positive controls (MAPF, taxol) and 6xHis-TTL3IDR, indicative of microtubule formation and tubulin stabilization, clear smearing of bands appears in the supernatant fractions of the negative controls (BSA, buffer), indicative no microtubule formation and tubulin degradation. M = Molecular marker; P = Pellet fraction; S = Supernatant fraction. (**G**) Identification of protein orthologs of Arabidopsis TTL1 and TTL3 in Embryophytes (earliest land plants), Chlorophyta, and Charophytes. The earliest TTL orthologs can only be identified in Embryophytes. IDR = intrinsically disordered region; TPR = tetratricopeptide repeat; TRX = C-terminal sequence with homology to thioredoxins and conserving the essential Cys residues required for thioredoxin activity; TRXL = TRX domain lacking essential Cys residues conserved for thioredoxin activity. Bootstrap test (0-100): 500 replicates; evolutionary distances: *p*-distance (0-1).

**Table S1.**
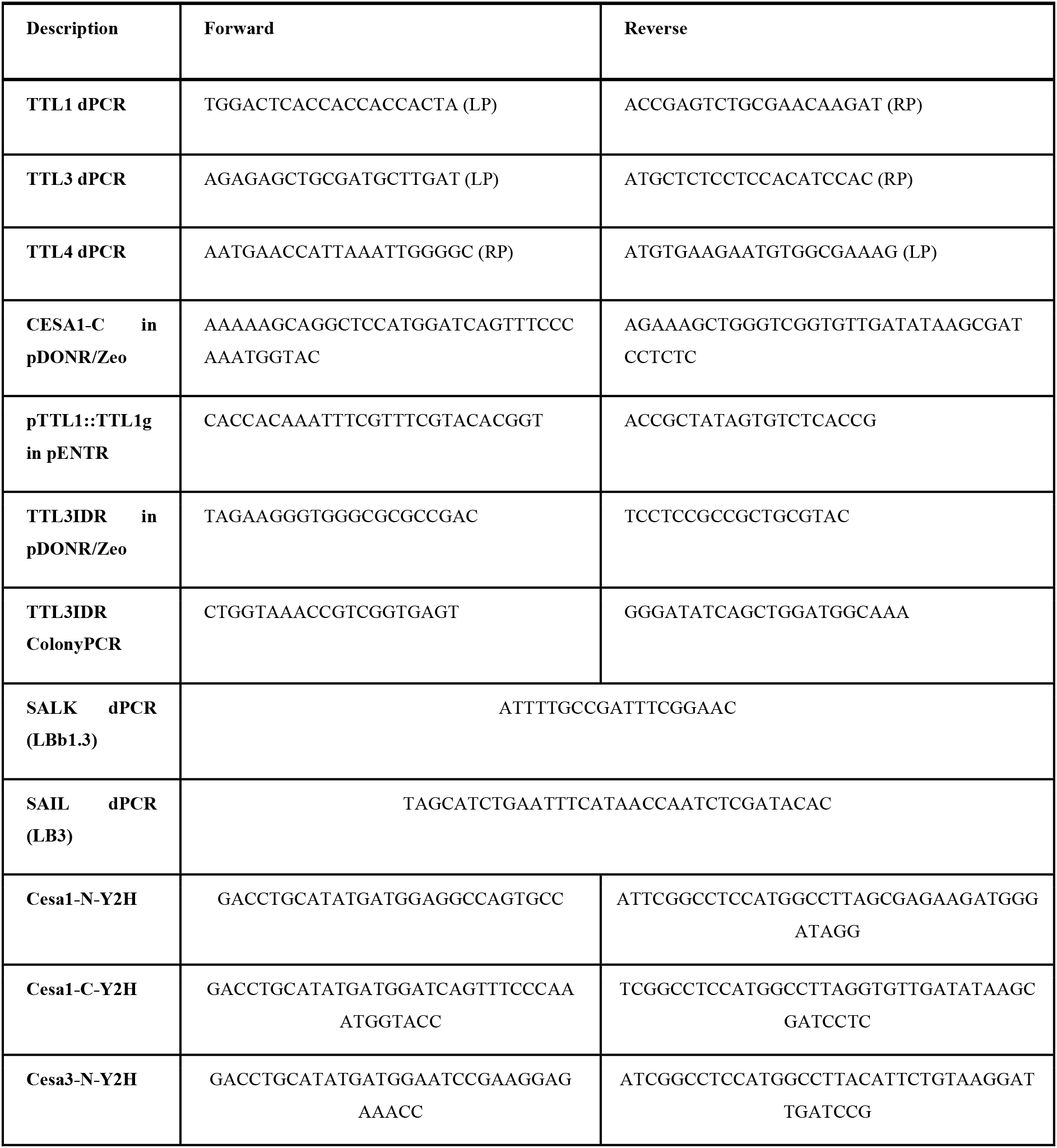
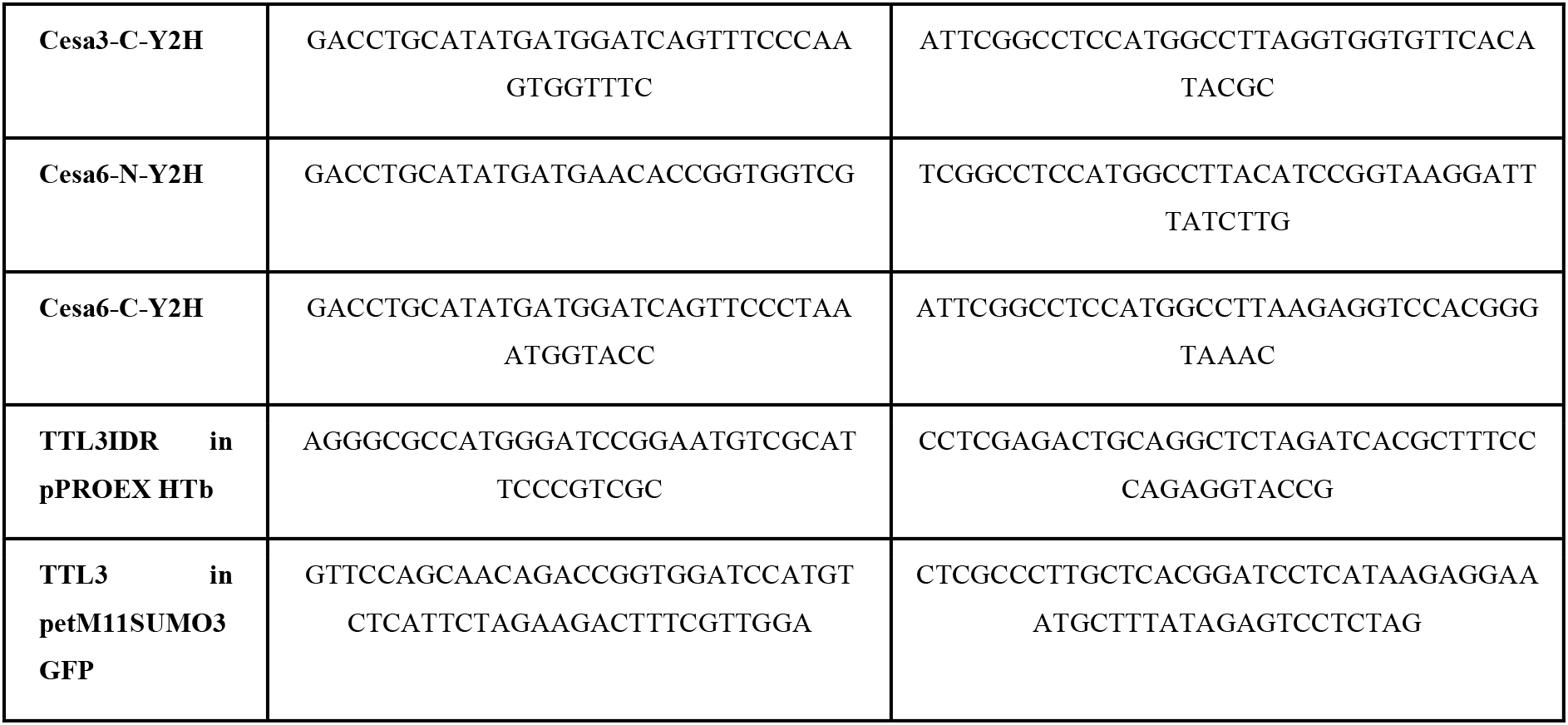
Primers used in this study.

## Caption for Movie S1

TTL3-GFP particles co-localize with tdT-CesA6 at the plasma membrane (see merged image series). TTL3-GFP image series are shown non-modified (non-mod) and after extracting particles at the plasma membrane with an image modification pipeline (mod).

